# An invariant C-terminal tryptophan in McdB mediates its interaction and positioning function with carboxysomes

**DOI:** 10.1101/2023.11.21.568049

**Authors:** Joseph L. Basalla, Maria Ghalmi, Y Hoang, Rachel Dow, Anthony G. Vecchiarelli

## Abstract

Bacterial microcompartments (BMCs) are widespread, protein-based organelles that regulate metabolism. The model for studying BMCs is the carboxysome, which facilitates carbon-fixation in several autotrophic bacteria. Carboxysomes can be distinguished as type α or ß, which are structurally and phyletically distinct. We recently characterized the Maintenance of Carboxysome Distribution (Mcd) systems responsible for spatially regulating α- and ß-carboxysomes, consisting of the proteins McdA and McdB. McdA is an ATPase that drives carboxysome positioning, and McdB is the adaptor protein that directly interacts with carboxysomes to provide cargo specificity. The molecular features of McdB proteins that specify their interactions with carboxysomes, and whether these are similar between α- and ß-carboxysomes, remain unknown. Here, we identify C-terminal motifs containing an invariant tryptophan necessary for α- and ß-McdBs to associate with α- and ß-carboxysomes, respectively. Substituting this tryptophan with other aromatic residues reveals corresponding gradients of carboxysome colocalization and positioning by McdB *in vivo*. Intriguingly, these gradients also correlate with the ability of McdB to form condensates *in vitro*. The results reveal a shared mechanism underlying McdB adaptor protein binding to carboxysomes, and potentially other BMCs. Our findings also implicate condensate formation as playing a key role in this association.

**SIGNIFICANCE STATEMENT:**

- Maintenance of carboxysome distribution protein B (McdB) is necessary for positioning a widespread class of protein-based organelles in bacteria that regulate metabolism. Without McdB, these organelles aggregate and lose functionality. How McdB interacts with and positions these organelles is unknown.
- We determine that an invariant tryptophan is necessary for McdB to interact with and position its organelle. A similar mechanism occurs in two diverse bacterial cell types, both relying on the invariant tryptophan.
- This class of bacterial organelle includes compartments involved in bacterial pathogenesis and carbon fixation. Our results therefore advance our understanding and applications of these organelles.

## INTRODUCTION

An important cellular feature across all domains of life is the compartmentalization of biological processes. Many bacteria possess protein-based organelles called bacterial microcompartments (BMCs) that provide subcellular compartmentalization and reaction isolation (1, 2). BMCs consist of selectively-permeable protein shells that encapsulate a set of enzymes, thus serving as nanoscale reaction centers for key metabolic steps (2). A recent bioinformatic survey identified 68 unique BMC types in 45 bacterial phyla (2), revealing that BMCs are widespread. Despite BMC prevalence and importance in diverse bacterial metabolisms, little is known about how BMCs are spatially regulated in the cell.

The best studied BMC type is the carboxysome (3). Carboxysomes encapsulate the enzyme ribulose-1,5-bisphosphate carboxylase/oxygenase (Rubisco) and co-concentrate it with its substrate CO_2_ to significantly increase the efficiency of carbon fixation in many autotrophic bacteria (3). As a result, carboxysomes are estimated to facilitate about 35% of all global carbon-fixation (4, 5), making carboxysomes of interest for developing carbon-capturing technologies (6, 7). Beyond their biotechnological potential, carboxysomes are also the paradigm for understanding fundamental aspects of general BMC biology, such as assembly, structure, and spatial regulation (1, 2, 3). Therefore, investigating the fundamental aspects of carboxysomes is important for developing technologies as well as deepening our understanding of BMCs.

Two subtypes of carboxysomes exist, α and ß, where ß-carboxysomes are found in ß-cyanobacteria and α-carboxysomes are found in numerous phylogenetically distinct groups, including α-cyanobacteria and several types of chemoautotrophic bacteria (2). While functionally equivalent, α- and ß-carboxysomes are structurally and phyletically distinct, with key differences in composition, mode of assembly, and regulation (8, 9). In fact, α-carboxysomes are more closely related to other BMC-types than they are to β-carboxysomes (9). Therefore, α- and ß-carboxysomes represent distinct BMC types, and comparative studies between the two have been critical for our understanding of carboxysome biology and BMCs in general.

Carboxysomes are spatially organized in the cell. In the model cyanobacterium *Synechococcus elongatus* PCC 7942 (*Se*, hereafter), a two-protein system is responsible for distributing ß-carboxysomes down the cell length (10, 11, 12). One component, which we named Maintenance of carboxysome distribution protein A (McdA), is a member of the ParA/MinD family of ATPases known to position various genetic and protein-based structures in bacteria (13, 14). The second component is a novel protein we named McdB, which interacts with McdA and also localizes to carboxysomes (10), thus acting as an adaptor to link the carboxysome cargo to its positioning ATPase (15). Deletion of either McdA or McdB results in carboxysome aggregation and asymmetric inheritance of carboxysome clusters, slower cell growth, and a rapid loss of carboxysomes in the cell population (16, 17). Therefore, uniform positioning maintains the carbon fixation efficiency of carboxysomes and ensures faithful inheritance of this vital organelle after cell division.

McdAB systems are widespread among β-cyanobacteria which contain β-carboxysomes, and proteobacteria which contain α-carboxysomes (11, 12). Using the α-carboxysome model organism *Halothiobacillus neapolitanus* (*Hn*, hereafter), we have shown that an McdAB system, distinct from that of β-carboxysomes, spatially distributes α-carboxysomes (12). Therefore, McdAB is a cross-phylum two-protein system necessary for positioning both α- and β-carboxysomes. More broadly, putative McdAB systems were also identified for other BMCs involved in diverse metabolic processes. Understanding how the McdAB system spatially regulates carboxysomes therefore has broad implications for understanding BMC trafficking across bacteria.

One outstanding question is how the adaptor protein, McdB, connects to and provides specificity for the carboxysome cargo. Our previous studies of β-McdB from *Se* and α-McdB from *Hn* revealed extreme differences at the sequence and structural levels (11, 12). For example, *Se* McdB is largely α-helical with a coiled-coil domain and forms a trimer-of-dimers hexamer (18), whereas *Hn* McdB is monomeric and completely intrinsically disordered (12). Therefore, given the extreme diversity between α- and β-McdB proteins, it also remains to be determined whether they follow a similar mechanism to associate with α- and ß-carboxysomes. Despite their diversity, α- and β-McdB proteins have been shown to form condensates *in vitro* (11, 12, 18). The processes underlying condensate formation *in vitro* can influence subcellular organization *in vivo* in both eukaryotes and prokaryotes (19, 20). Along these lines, we recently found evidence suggesting that condensate formation by *Se* McdB may play a role in its association with carboxysomes *in vivo* (21). To what extent condensate formation by McdB influences its association with carboxysomes, and whether this activity plays a role in the spatial organization of α- and ß-carboxysomes remains to be determined.

Here, we identified a C-terminal motif that contains an invariant tryptophan in both α - and β-McdB proteins. We determined this invariant tryptophan is essential for the association of McdB with β-carboxysomes in *Se* and α-carboxysomes in *Hn*. Furthermore, expressing only this C-terminal motif containing the tryptophan and surrounding residues was necessary and sufficient for carboxysome association in *Hn*, but not in *Se*. We provide evidence to suggest *Se* McdB oligomerization is also required. We also show that putative McdB-like proteins associated with other BMC types encode invariant tyrosines, suggesting other aromatic residues can serve the same role as tryptophan (12). By substituting the C-terminal tryptophan in *Hn* and *Se* McdB with other aromatic residues, we observed corresponding complementation gradients of carboxysome association and carboxysome positioning. Interestingly, these gradients of activity *in vivo* correlated with condensate formation *in vitro* for the purified aromatic mutants of McdB. Together, the results show that despite the extreme diversity between α- and β-McdB proteins and α- and β-carboxysomes, a similar mode of association is used. These results lay the groundwork for understanding the molecular mechanisms of protein association with the surface of the carboxysome and potentially other BMCs across bacteria.

## RESULTS

### All McdB proteins encode an invariant tryptophan at their extreme C-terminus

We first performed multiple sequences alignments both within and across α- and ß-McdB types to identify regions of conservation that may be involved in associating with carboxysomes. On average, McdBs show low sequence identity; 14.8% among α-McdBs (Figure 1A), 14.9% among ß-McdBs (Figure 1B), and 6.7% across all McdBs (Figure 1C). One reason for this low average identity is the large alignment gaps that stem from the high variance in sequence lengths, which range from 51 – 169 residues for α-McdBs and 132 – 394 residues for ß-McdBs (Supplemental Figure S1A-C). Although the average identity was low, we identified three invariant residues (WPD) at the C-terminus of α-McdBs (Figure 1A), and one invariant residue (W) at the C-terminus of ß-McdBs (Figure 1B), consistent with our previous reports (11, 12). Aligning all full-length McdB protein sequences did not identify any invariant residues (Figure 1C), again due to the large variations in protein lengths, which prevented the C-termini of α- and ß-McdBs from aligning (Supplemental Figure S1C). We therefore repeated these alignments on only the last 20 C-terminal amino acids of all McdB sequences to control for length variation (Figure 1D-F). These alignments unveiled motif conservation specific to α - and ß-McdB types as well as conservation shared across all McdBs. α-McdBs contain a short consensus sequence of R(V/I)WPD at the extreme C-terminus (Figure 1D). The ß-McdB C-terminal motif is more degenerate, showing an enrichment of aspartic acids (D) within the last 10 C-terminal residues, along with the invariant tryptophan (Figure 1E). And across all McdBs, we identified the tryptophan as the only invariant residue (Figure 1F), suggesting a critical role in its functionality for both α- and ß-McdBs.

**Figure 1:**
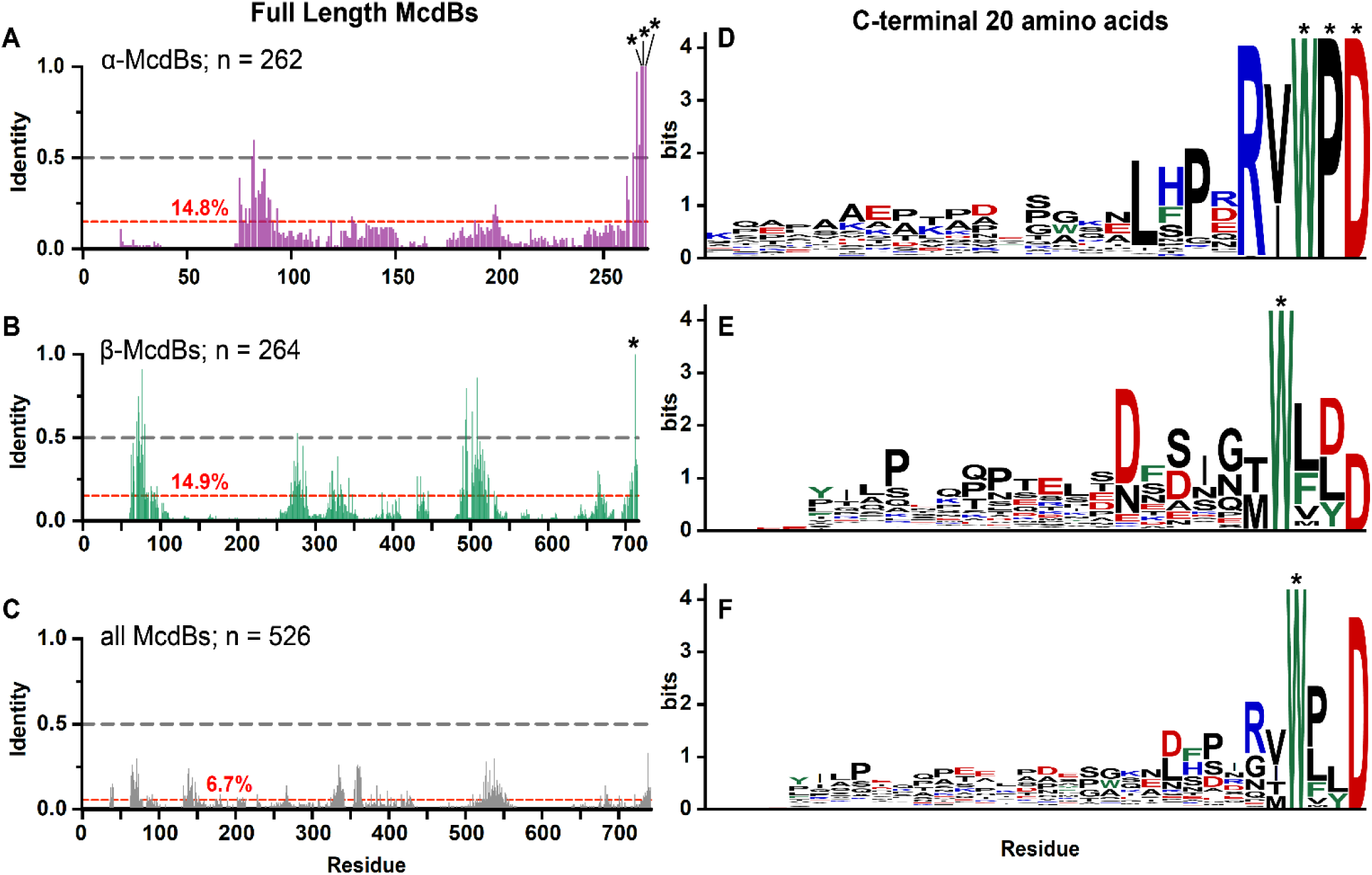
All McdB proteins share an invariant C-terminal tryptophan. (**A-C**) Percent identities from multiple sequence alignments of full-length (**A**) α-McdBs, (**B**) ß-McdBs, and (**C**) both α- and ß-McdBs. The average percent identity for each alignment is shown in red. (**D-F**) Sequence logos generated from multiple sequence alignments of only the last 20 C-terminal amino acids of (**D**) α-McdBs, (**E**) ß-McdBs, and (**F)** both α- and ß-McdBs. Positions that are invariant (100 percent identity) are indicated with an asterisk. Cationic residues are colored blue, anionic residues red, aromatic residues green, and all others black.

### The C-terminal invariant tryptophan is required for McdB association with carboxysomes

Adaptor proteins typically associate with their cognate ParA/MinD positioning ATPase via basic residues in the N-terminus of the adaptor (22), and we have computationally shown that this is likely the case for McdB interactions with McdA (23). Therefore, our identification of an invariant tryptophan within the C-terminus of all McdBs motivated our study for its potential role in associating with the carboxysome cargo, as opposed to McdA, which we expect McdB interacts with via its N-terminus.

We performed *in vivo* fluorescence microscopy in both *Se* and *Hn* cells to determine how McdB localization and carboxysome organization were altered for McdB mutants lacking the invariant C-terminal tryptophan. To visualize carboxysomes, the fluorescent protein monomeric Turquoise2 (mTQ) (24) was fused to the C-terminus of the small subunit of the Rubisco enzyme (RbcS in *Se*; CbbS in *Hn*). RbcS-mTQ and CbbS-mTQ were expressed using a second copy of their native promoters inserted at a neutral site, in addition to the wild type copy at the native locus. To simultaneously image McdB mutants in these carboxysome reporter strains, mutations were made in an McdB variant that was N-terminally fused to the fluorescent protein monomeric NeonGreen (mNG) (25). We have previously shown that, in *Se*, mNG-McdB is fully functional for carboxysome positioning when expressed as the only copy of McdB at its native locus (10). In *Hn*, the mNG fusion unfortunately perturbs McdB interactions with McdA, resulting in carboxysome aggregation. However, this fusion still associates with carboxysomes and therefore remains a useful positive control for studying McdB-carboxysome association in *Hn* (12). Finally, we also performed phase-contrast imaging to monitor cell morphology.

In *Se*, the invariant tryptophan is the final C-terminal amino acid (Supplemental Figure S1D), therefore we simply deleted it to make McdB[ΔW152]. As shown previously in wild type *Se* cells (10), mNG-McdB colocalized with carboxysome foci that are uniformly distributed down the cell length (Figure 2A). Without the invariant tryptophan, mNG-McdB[ΔW152] was diffuse in the cell (Figure 2A), with no notable carboxysome colocalization compared to that of wild type (Figure 2B). Carboxysomes were also mispositioned (Figure 2C) and clustered into high-intensity aggregates in the McdB[ΔW152] strain (Supplementary Figure S2B), which phenocopies a complete deletion of McdB, as shown previously (17).

**Figure 2:**
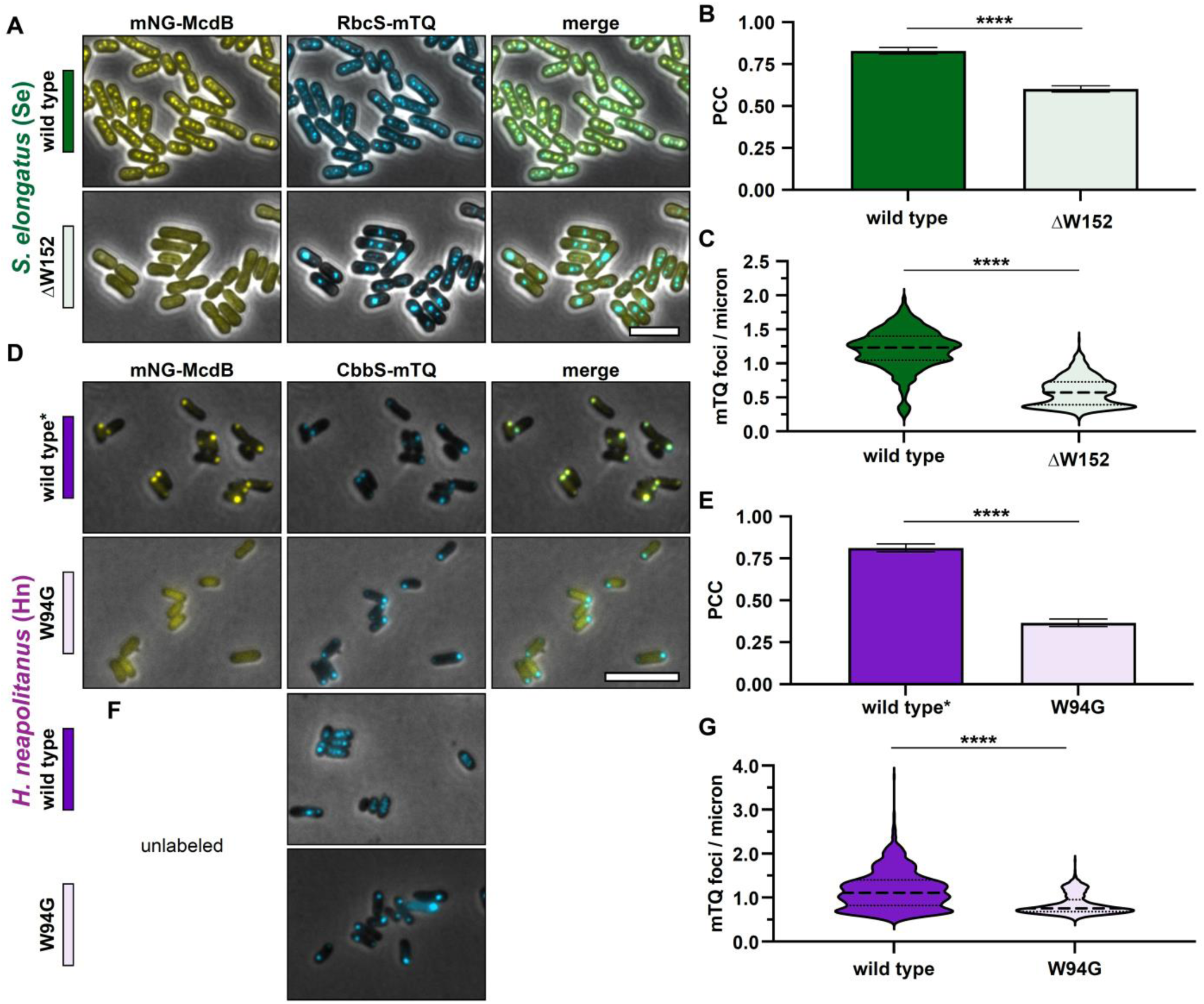
The invariant tryptophan for both α- and β-McdBs mediates carboxysome localization. (**A**) Representative microscopy images of the indicated *Synechococcus elongatus* (*Se*) strains. Phase contrast images are shown in black and white and overlaid with the fluorescence channels: mNG-McdB proteins are yellow and RbcS-mTQ labelled carboxysomes are cyan. Colored bars next to the strain names correspond to colors on the associated graphs. (**B**) Pearson’s Correlation Coefficients (PCCs) quantified for the indicated *Se* strains. Graphs represent means and standard deviations from 7 technical replicates each with n > 500 cells. **** p < 0.001 from Welch’s t-test. (**C**) Quantification of carboxysome spacing as number of mTQ foci divided by cell length. Graphs represent medians and interquartile ranges from 3 biological replicates each with n > 500 cells. **** p < 0.001 from Mann-Whitney U-test. (**D**) Representative microscopy images of the indicated *Halothiobacillus neapolitanus* (*Hn*) strains. Wild type* indicates the wild type McdB with an N-terminal mNG tag, which causes carboxysome aggregation in *Hn*. Phase contrast images are overlaid with the fluorescence channels: mNG-McdB proteins are yellow and CbbS-mTQ labelled carboxysomes are cyan. Colored bars next to the strain names correspond to colors on the associated graphs. (**E**). Pearson’s Correlation Coefficients (PCCs) quantified for the indicated *Hn* strains. Graphs represent means and standard deviations from 7 technical replicates each with n > 500 cells. **** p < 0.001 from Welch’s t-test. (**F**) As in (D), but with McdB not labeled with mNG. (**G**) Quantification of carboxysome spacing as number of mTQ foci divided by cell length. Graphs represent medians and interquartile ranges from 3 biological replicates each with n > 500 cells. **** p < 0.001 from Mann-Whitney U-test. Scale bars are 5 µm and apply to all images.

For *Hn* McdB, the invariant tryptophan is the third amino acid from the C-terminus (Supplemental Figure S1D). Therefore, a glycine substitution was used to make McdB[W94G]. Although the mNG-McdB fusion destroys its carboxysome positioning function, the fusion still strongly colocalized with the mispositioned carboxysome aggregates at the cell pole, hence the label “wild type*” (Figure 2D). As with *Se* McdB[ΔW152], mNG-McdB[W94G] was diffuse in *Hn* cells and did not colocalize with carboxysomes (Figure 2E).

While mNG-McdB[W94G] allowed us to observe changes in protein localization, we were unable to determine the effects of this mutant on carboxysome positioning when fused to mNG. Therefore, we also imaged carboxysome distribution in *Hn* cells with unlabeled McdB[W94G] (Figure 2F). Consistent with *Se* McdB[ΔW152], carboxysomes became mispositioned (Figure 2G) and clustered into high-intensity aggregates in the McdB[W94G] mutant (Supplementary Figure S2B), which phenocopies a complete deletion of McdB in *Hn*, as shown previously (12). Together, our results show that the C-terminal tryptophan found in all McdB proteins is necessary for carboxysome association and positioning.

### Loss of carboxysome association and positioning is not due to destabilization of McdB mutants

We found it striking that a single residue change completely destroyed McdB association and positioning in both *Se* and *Hn* cells. We therefore set out to confirm that the observed phenotypes were not a consequence of these mutations destabilizing McdB. We used circular dichroism (CD) and size-exclusion chromatography with multi-angle light scattering (SEC-MALS) to determine the secondary and quaternary structures of the purified proteins, respectively. As shown previously (12, 18), wild type *Se* McdB forms a hexamer in solution with an α-helical signature, whereas *Hn* McdB is monomeric and completely intrinsically disordered (Supplemental Figure S3A-B). Since wild type *Hn* McdB is monomeric and disordered, there is no structure to disrupt in McdB[W94G]. We therefore focused our analyses on McdB[ΔW152]. *Se* McdB[ΔW152] remained hexameric and displayed the same α-helical signature as that of wild type *Se* McdB, showing this mutation did not destabilize *Se* McdB structure or oligomerization *in vitro*.

Furthermore, we then confirmed that the tryptophan deletion did not destabilize *Se* McdB structure *in vivo*, potentially leading to degradation. The average mNG fluorescence per cell was quantified and compared between *Se* strains with wild type McdB and McdB[ΔW152]. No decrease in the average mNG signal was observed in the McdB[ΔW152] strain, compared to that of wild type (Supplemental Figure S3C). Together, the results show that the tryptophan mutations have no significant effects on McdB stability either *in vivo* or *in vitro,* and indicate that the invariant tryptophan directly mediates α- and ß-McdB interactions with their respective carboxysomes.

### The C-terminus of monomeric Hn McdB is necessary and sufficient for carboxysome association, but not for hexameric Se McdB

We next set out to determine if the conserved C-terminal motifs we identified, which contain the invariant tryptophan, were sufficient to drive McdB association with carboxysomes. To investigate this, we N-terminally fused mNG to the last 31 amino acids of *Se* McdB and the last 10 amino acids of *Hn* McdB (Figure 3A). The choice of C-terminal domain (CTD) size was informed by our previous biochemical analysis of *Se* McdB, which revealed folded regions extending ∼ 30 amino acids from the C-terminus (18). We aimed to preserve this folding in the event that it was required for *Se* McdB association with carboxysomes. *Hn* McdB, on the other hand, is monomeric and completely disordered (Supplemental Figure S3A-B). Therefore for the *Hn* McdB[CTD] construct, only the last 10 amino acids were included, comprising the highest identities from the α-McdB C-terminal alignments (see Figure 1D).

**Figure 3:**
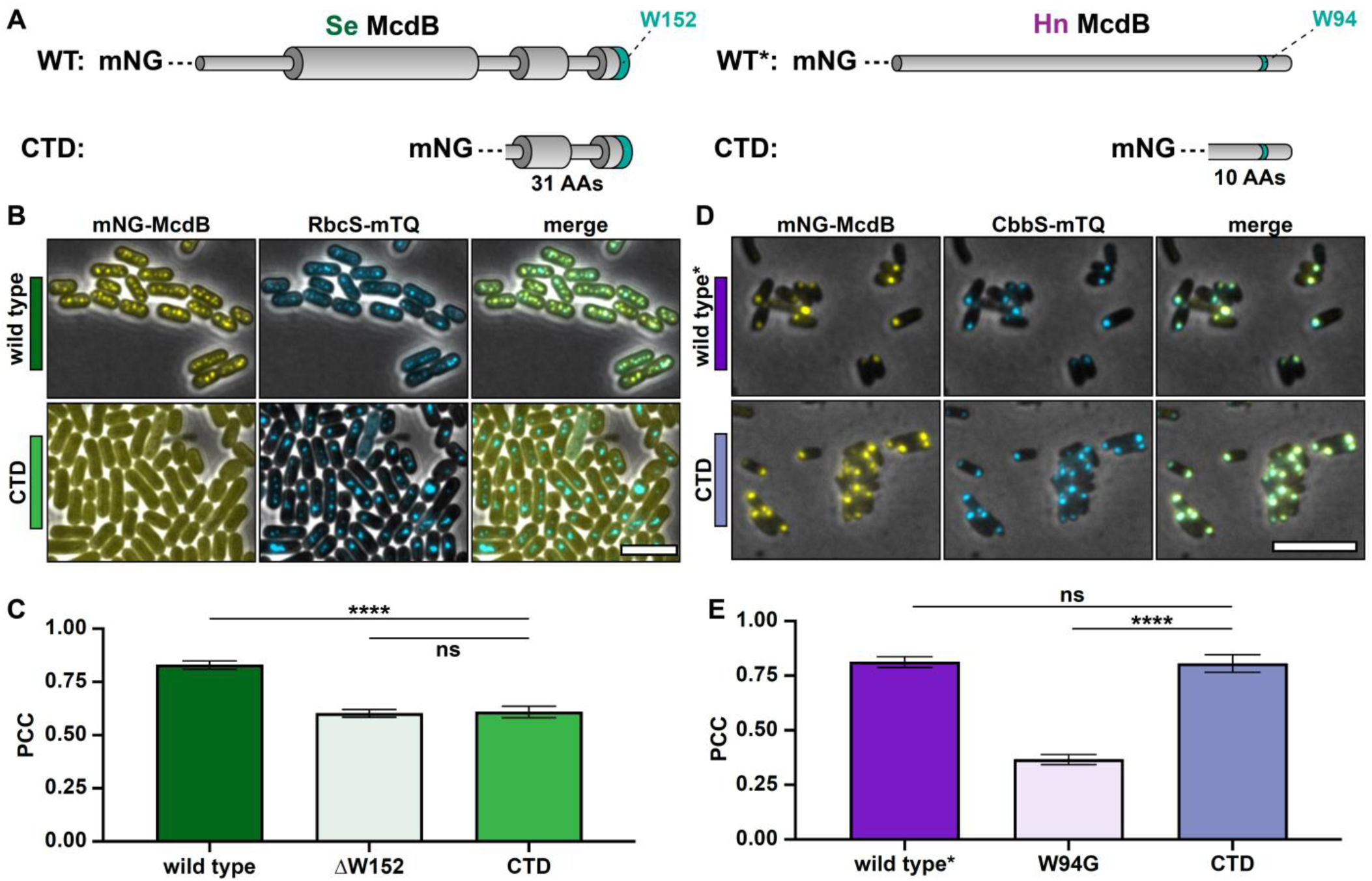
The C-termini of α- and β-McdBs show differences in their ability to localize to carboxysomes. (**A**) McdB protein models for *Se* and *Hn* wild type (WT) and C-terminal domains (CTD). Wide cylinders represent α-helical regions and narrow cylinders represent region of intrinsic disorder. The invariant tryptophan (W) is represented as a green stripe in the protein models. Sizes of the CTD truncations used are indicated below the respective model. (**B**) Representative microscopy images of the indicated *Se* strains. Phase contrast images are shown in black and white and overlaid with the fluorescence channels: mNG-McdB proteins are yellow and RbcS-mTQ labelled carboxysomes are cyan. Colored bars next to the strain names correspond to colors on the associated graphs. (**C**). Pearson’s Correlation Coefficients (PCCs) quantified for the indicated *Se* strains. Graphs represent means and standard deviations from 7 technical replicates each with n > 500 cells. **** p < 0.001 and ns = non-significant from Welch’s t-test. (**D**) Representative microscopy images of the indicated *Hn* strains. Wild type* indicates the wild type McdB with an N-terminal mNG tag, which causes carboxysome aggregation in *Hn*. Phase contrast images are overlaid with the fluorescence channels: mNG-McdB proteins are yellow and CbbS-mTQ labelled carboxysomes are cyan. Colored bars next to the strain names correspond to colors on the associated graphs. (**E**) Pearson’s Correlation Coefficients (PCCs) quantified for the indicated *Se* strains. Graphs represent means and standard deviations from 7 technical replicates each with n > 500 cells. **** p < 0.001 and ns = non-significant from Welch’s t-test. Scale bars are 5 µm and apply to all images.

Intriguingly, the mNG-CTDs from *Se* and *Hn* McdB displayed different localizations. *Se* McdB[CTD] was completely diffuse in the cell (Figure 3B), and showed no association with carboxysomes, similar to that of McdB[ΔW152] (Figure 3B-C). *Hn* McdB[CTD], on the other hand, strongly colocalized with carboxysomes (Figure 3D), similar to that of wildtype* (Figure 3E). The data show that the last 10 amino acids of *Hn* McdB are necessary and sufficient for associating with α-carboxysomes. However, the last 31 amino acids of *Se* McdB, despite encoding the conserved motif and invariant tryptophan, is insufficient.

Recall that full-length *Se* McdB is a hexamer in solution whereas *Hn* McdB is monomeric (Supplemental Figure S3A). Furthermore, we have previously shown that the CTD of *Se* McdB has an α-helical secondary structure (18), while *Hn* McdB is completely disordered (12). It is therefore possible that the *Se* McdB[CTD] has altered protein structure and/or oligomerization that influences its association with carboxysomes. To investigate oligomerization, we performed SEC on *Se* McdB[CTD] (3.7 KDa), and used full-length *Hn* McdB (10 KDa) and an N-terminal peptide of *Se* McdB (*Se* McdB[NTD]; 2.3 kDa) as sizing standards. Both *Se* McdB[CTD] and *Se* McdB[NTD] eluted at the lower end of the separation range of the column (3 kDa) (Supplemental Figure S4A), indicating that *Se* McdB[CTD] remains monomeric. CD analysis confirmed that *Se* McdB[CTD] retained an α-helical structure (Supplemental Figure S4B). Full-length *Hn* McdB and *Se* McdB[NTD] are provided as disordered protein controls. The retained α-helical structure of *Se* McdB[CTD] explains why it eluted later than the disordered *Se* McdB[NTD] construct, despite having a higher molecular weight (Supplemental Figure S4A).

Together, the results show that *Se* McdB[CTD] retains its α-helical structure, but does not form a hexamer like the full-length protein. We propose that the C-terminal association of *Se* McdB with carboxysomes requires higher avidity provided by hexamerization, whereas for the *Hn* McdB monomer, a single motif is necessary and sufficient for carboxysome association.

### Other aromatic residues functionally replace the invariant tryptophan with varying activity

Our previous bioinformatic analyses identified putative McdB-like proteins associated with other BMCs (11), including the 1,2-propanediol utilization microcompartment (PDU) and the glycyl radical enzyme-containing microcompartment (GRM) (1). Intriguingly, the C-termini of these McdB-like proteins lack the invariant tryptophan (Figure 4A). Instead, we found tyrosine (Y) or phenylalanine (F) residues conserved within the last five amino acids (Figure 4B), suggesting other C-terminal aromatics could potentially fulfill the role of the invariant tryptophan.

**Figure 4:**
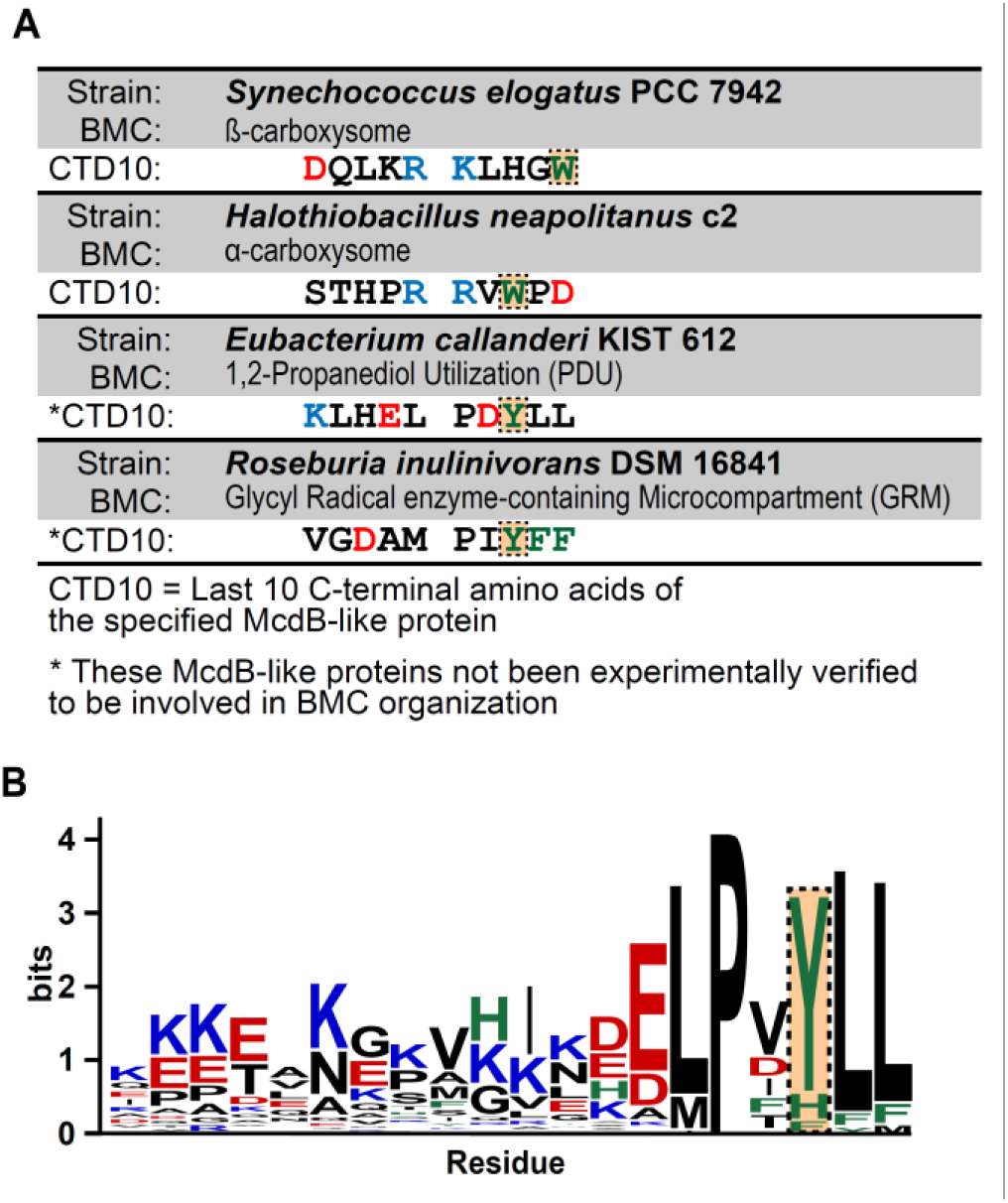
McdB-like proteins found near the operons of non-carboxysome BMCs have aromatics other than tryptophan at their C-termini. (**A**) Table displaying the last 10 C-terminal amino acids of McdBs from *Se* and *Hn*, as well as from putative McdBs from bacteria containing the indicated BMCs. Acidic residues are colored red, basic residues blue, and aromatic residues green. The residues at the position of the invariant tryptophan are boxed and highlighted. (**B**) Sequence logos generated from a multiple sequence alignments of the last 20 C-terminal amino acids of McdBs from non-carboxysome BMCs. The position corresponding to the invariant tryptophan as shown in (A) is boxed and highlighted.

To test this, we substituted the invariant tryptophan with Y or F in both the *Se* and *Hn* McdB proteins. These McdB variants were N-terminally fused to mNG, expressed at the native locus, and imaged in the carboxysome-labeled strains. We found that *Se* McdB[W152Y] colocalized with carboxysomes to the same degree as wild type McdB (Figure 5A-B). *Se* McdB[W152F] showed weaker association, but still greater than *Se* McdB[ΔW152]. Intriguingly, this gradient of carboxysome colocalization strongly correlated with the carboxysome positioning function of each *Se* McdB variant (Figure 5C, Supplemental Figure S5A). *Se* McdB[W152Y] still distributed carboxysomes, albeit with slightly perturbed spacing and higher foci intensities compared to that of wild type. *Se* McdB[W152F] showed an even lesser degree of carboxysome positioning compared to McdB[W152Y], but still greater than *Se* McdB[ΔW152].

**Figure 5:**
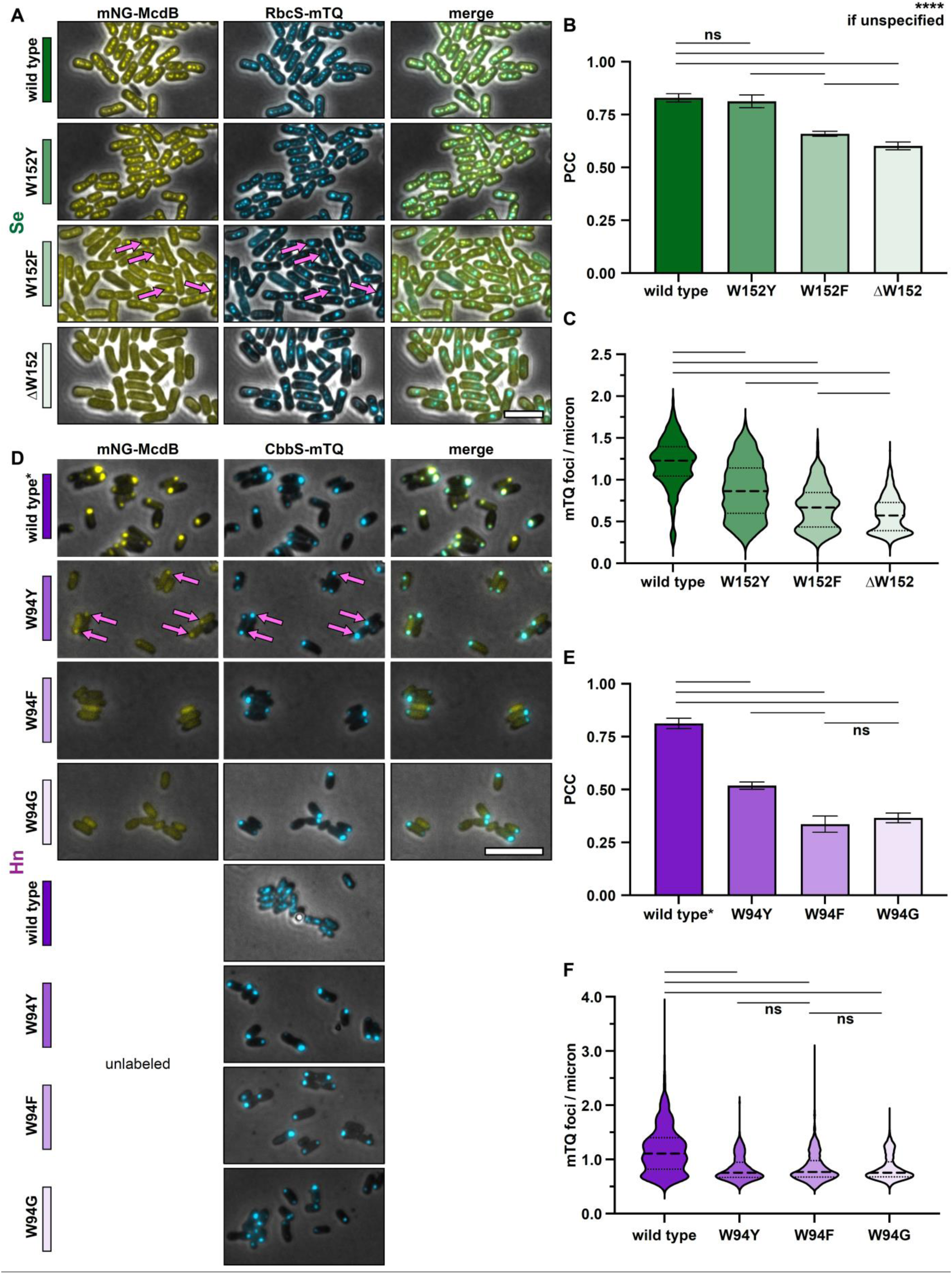
Changing the invariant tryptophan to other aromatic residues reveals a gradient of McdB colocalization with carboxysomes. (**A**) Representative microscopy images of the indicated *Se* strains. Phase contrast images are shown in black and white and overlaid with the fluorescence channels: mNG-McdB proteins are yellow and RbcS-mTQ labelled carboxysomes are cyan. Magenta arrows highlight moderate McdB colocalization with carboxysomes. Colored bars next to the strain names correspond to colors on the associated graphs. (**B**). Pearson’s Correlation Coefficients (PCCs) quantified for the indicated *Se* strains. Graphs represent means and standard deviations from 7 technical replicates each with n > 500 cells. **** p < 0.001 and ns = non-significant from Welch’s t-test. (**C**) Quantification of carboxysome spacing as number of mTQ foci divided by cell length. Graphs represent medians and interquartile ranges from 3 biological replicates each with n > 500 cells. **** p < 0.001 from Mann-Whitney U-test. (**D-F**) As in (A-C), but in *Hn* strains. “Unlabeled” refers to the strain set with the indicated mutations in McdB, but McdB is not labeled with mNG. Wild type* indicates the wild type McdB with an N-terminal mNG tag, which causes carboxysome aggregation in *Hn*. Scale bars are 5 µm and apply to all images.

In *Hn*, substituting the tryptophan with other aromatic residues was significantly less permissive. McdB[W94Y] only moderately colocalized with carboxysomes, whereas McdB[W94F] was completely diffuse in the cell, similar to that of McdB[W94G] (Figure 5E). And none of the McdB mutants were capable of positioning carboxysomes in *Hn* (Figure 5F, Supplemental Figure S5B).

To summarize, we found a striking gradient of McdB variant colocalization with carboxysomes that followed similar trends: W152 ≈ W152Y > W152F > ΔW152 in *Se* (Figure 5B), and W94 > W94Y > W94F ≈ W94G in *Hn* (Figure 5E). Furthermore, in *Se*, the gradient of carboxysome association directly correlated with the carboxysome positioning function of the McdB mutants (Figure 5C, Supplemental Figure S5A). In *Hn*, however, none of the McdB mutants restored carboxysome positioning (Figure 5F, Supplemental Figure S5B).

Overall, in both *Se* and *Hn*, substituting the conserved tryptophan with tyrosine provided strong McdB localization to carboxysomes, compared to that of the phenylalanine substitution. In fact, *Se* McdB[W152Y] localized to carboxysomes to a comparable degree as wild type McdB and provided near-wild type carboxysome positioning function. This is striking given that all carboxysome-associated McdBs bioinformatically identified encode an invariant C-terminal W instead of Y, and W is generally considered the least substitutable amino acid (26). We conclude that C-terminal aromatic residues can drive the localization of McdBs to both α- and ß-carboxysomes, and suggests a similar role for the conserved C-terminal aromatic residues found in putative McdB-like proteins associated with other BMC types.

### The invariant tryptophan influences McdB condensate formation

We previously found that several McdB proteins, including those from *Se* and *Hn*, can form biomolecular condensates *in vitro* (11, 12, 18, 20). For *Se* McdB, positively charged residues within the disordered N-terminus mediate the degree to which McdB forms condensates - as positive charges were removed, the ability to form condensates decreased (21). Our biochemical characterization also suggested these N-terminal positive residues may associate with the C-terminus of other McdB molecules to drive condensation. However, we did not identify C-terminal McdB mutants that altered condensation without destroying hexamerization.

Some protein condensates form via cation-π networks, where positively charged residues interact with electron-dense aromatic residues through electrons in their π orbitals (27, 28). Interestingly, these studies have shown gradients of condensate formation by changing the type of aromatic residues involved (27). We set out to determine if the invariant tryptophan influenced McdB condensate formation *in vitro*, and whether other aromatic residues at this position resulted in a gradient of condensate forming activity that could provide mechanistic insight into our observations *in vivo*.

We purified the *Se* McdB variants and compared the degree to which each formed condensates under conditions we previously found to facilitate condensate formation for wild type McdB (18). The level of condensate formation for McdB[W152Y] was slightly lower than wild type, and lower still for McdB[W152F]. *Se* McdB[ΔW152] could form condensates, but with the lowest activity (Figure 6A). Intriguingly, this gradient of condensation activity (W152 > W152Y > W152F > ΔW152) directly correlates with the ability of these *Se* McdB variants to associate with carboxysomes and drive their positioning reactions *in vivo* (See Figure 5A-C).

**Figure 6:**
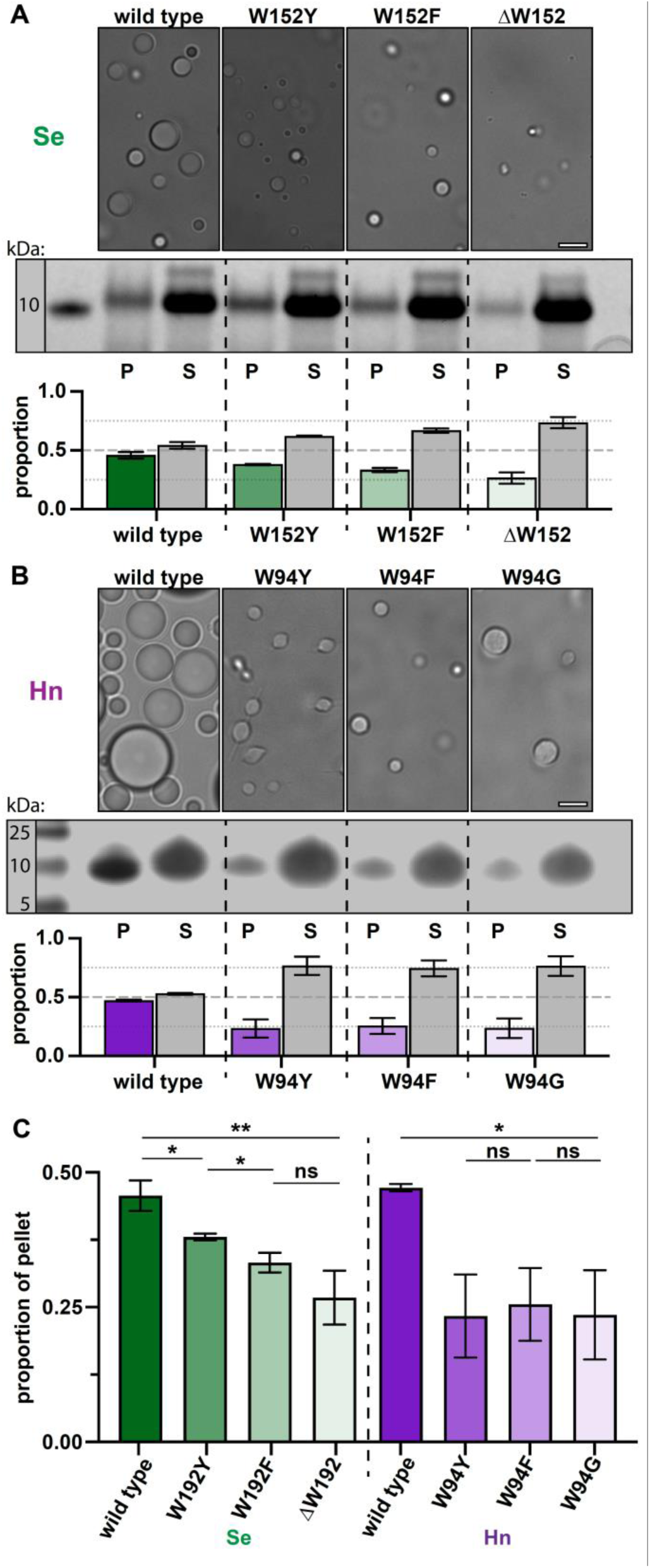
Changing the invariant tryptophan to other aromatic residues reveals a gradient of condensate formation for *Se* McdB but not *Hn* McdB. (**A**) (*top*) Representative DIC microscopy images of the indicated *Se* McdB variants at 50 µM (100 mM KCl; 20 mM HEPES, pH 7.2) after 1 hour. (*middle*) Samples were pelleted (P) and run on SDS-PAGE gel along with associated supernatant (S). (*bottom*) Gel bands were quantified. Graphs represent the proportion of total intensity from a P / S pair, and are reported as the mean and standard deviation from 3 replicates. (**B**) As in (A), but for *Hn* McdB at 700 µM (100 mM KCl; 20 mM HEPES, pH 7.2; 15% PEG-8000) after 18 hours. (**C**) Quantification summary of pellet fractions from *Se*- and *Hn*-McdB variants. ** p < 0.01, * p < 0.05, ns = non-significant from Welch’s t-test. Scale bars are 5 µm and apply to all microscopy images.

When performing the same comparison with the purified McdB variants from *Hn*, we did not observe a gradient of condensate formation as we did with the *Se* proteins. Instead, all *Hn* McdB mutants showed the same significant loss of condensate formation (Figure 6B-C), which correlates with the inability of these *Hn* McdB variants to associate with and position carboxysomes *in vivo* (See Figure 5D-F).

Finally, to investigate the condensate forming activity and relative expression levels of McdB variants *in vivo*, we expressed mCherry-tagged McdB constructs in *E. coli* and monitored the formation of foci as well as expression levels over time. Using this approach, we have recently shown that the fluorescent foci formed by *Se* mCherry-McdB in *E.coli* cells (Supplemental Figure S6A) are liquid-like condensates (18, 29). Intriguingly, the tryptophan mutants once again displayed the same functional gradient in forming condensates *in vivo* (Supplemental Figure S6B). Importantly, all *Se* McdB variants showed the same levels of expression compared to wildtype, and with no notable degradation (Supplemental Figure S6C). The data provide an additional line of evidence showing that mutation to the invariant tryptophan did not result in the destabilization and/or degradation of McdB variants. Together, we once again find a gradient of condensation activity *in vivo* that mirrors our *in vitro* results with *Se* McdB (W152 > W152Y > W152F > ΔW152).

*Hn* McdB remained soluble in *E. coli*, even at the highest expression levels achievable in this assay (Supplemental Figure S6D). This is consistent with *Hn* McdB requiring significantly higher protein concentrations to form condensates *in vitro*, compared to *Se* McdB (see Figure 6A-B). Since we could not form foci with wildtype *Hn* McdB, mutant versions of *Hn* McdB were not pursued using this approach.

Overall, the data implicate the invariant C-terminal tryptophan as a major contributor to condensate formation for both α- and β-McdBs. Furthermore, aromatic residue substitutions at the tryptophan position can affect McdB condensate formation *in vitro* in a manner that directly correlates with how aromatics affect McdB function *in vivo*.

## DISCUSSION

Here, we identified and probed the function of conserved C-terminal motifs containing a tryptophan residue that is invariant across all McdB proteins bioinformatically identified to date. The invariance is striking because even when comparing small regions of McdB sequences of fixed length, this tryptophan is the only residue with 100% identity (Figure 1). We found that the invariant tryptophan is necessary for α- and ß-McdBs to colocalize with α- and ß-carboxysomes, respectively. With the invariant tryptophan removed, McdB became diffuse in the cytoplasm (Figure 2). Interestingly, the C-terminal motif containing the invariant tryptophan was necessary and sufficient for carboxysome localization by α-McdB of *Hn*, but not for ß-McdB of *Se* (Figure 3). We suggest this discrepancy may be due to differences in the minimal oligomeric unit of the proteins, whereby full-length *Hn* McdB and the C-terminal fragment are both monomers, while full-length *Se* McdB is a hexamer and its C-terminal fragment is monomeric (Supplemental Figure S4).

We also found that putative McdB-like proteins that are associated with other BMCs have conserved C-terminal aromatic residues other than tryptophan (Figure 4). We therefore attempted to complement the removal of the invariant tryptophan in both α- and ß-McdBs by substituting it with other aromatic amino acids. Intriguingly, we observed a gradient of carboxysome association that correlated with the carboxysome positioning function of each McdB variant (Figure 5). Lastly, this gradient of function *in vivo* directly correlated with the ability of these McdB variants to form condensates both *in vitro* (Figure 6) and *in vivo* (Supplemental Figure S6D). A summary of our findings is provided in Figure 7. Together, the data provide a foundation for future studies on the molecular nature of McdB-carboxysome interactions and the role protein condensation may play in this association.

**Figure 7:**
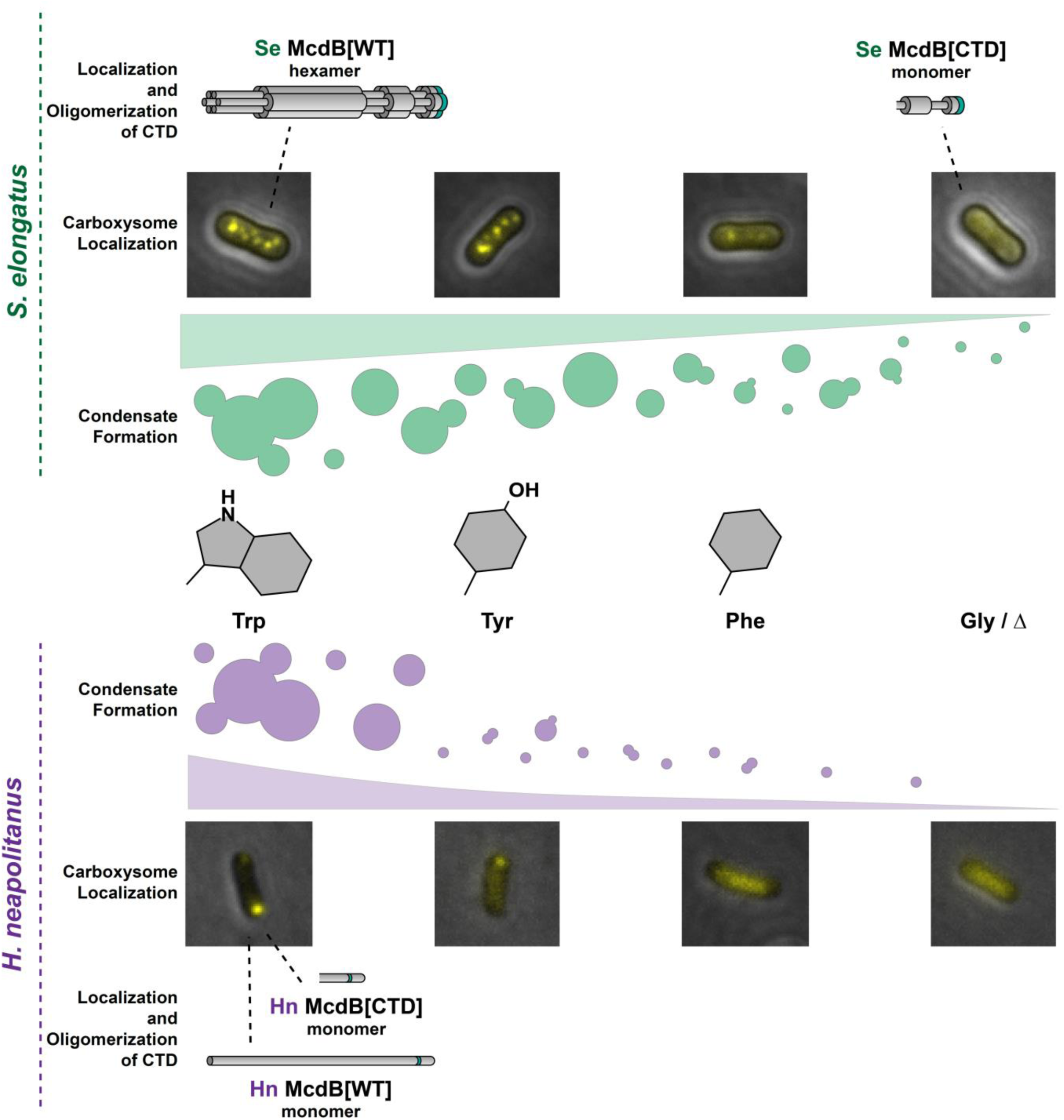
Summary of condensate formation and carboxysome localization for the aromatic substitution mutants of *Se* (*top*) and *Hn* (*bottom*) McdB. Wild type (WT) McdB proteins from *Se* and *Hn* have an invariant C-terminal tryptophan (trp), depicted as a green stripe in the cartoon protein models. *Se* McdB functions as a hexamer whereas *Hn* McdB is monomeric. We substituted this trp with tyrosine (tyr), phenylalanine (phe), glycine (gly) as well as deleted the trp (Δ) (*center*), which revealed a gradient of condensate formation activity and carboxysome localization (ramps). The C-terminal domain (CTD) containing the invariant tryptophan was sufficient to localize *Hn* McdB to carboxysomes (*bottom*). However this was not the case for *Se* McdB[CTD] suggesting oligomerization of *Se* McdB is also a requirement (*top*).

### Comparative analyses on BMC shell proteins could further our understanding of their molecular interactions with McdB proteins

It remains to be determined how, at a molecular level, the invariant tryptophan drives McdB association with carboxysomes. Carboxysomes, like all BMCs, are comprised of a selectively permeable protein shell and an enzymatic core (1). While the set of encapsulated core enzymes are highly diverse, the outer shell proteins of BMCs are well-conserved in sequence and structure (1, 2). For all BMCs, several different types of protein oligomers build the outer shell to give rise to the characteristic polyhedral shape of a BMC (1); the most abundant of which is a hexamer (BMC-H). For β-carboxysomes, the major hexameric shell protein is called CcmK2. In *Se*, we have shown that McdB strongly associates with CcmK2 (10). However, the regions and residues of CcmK2 required for the McdB-CcmK2 association remain to be determined.

Although BMC-H proteins have regions of high conservation, there are also variable regions that have led to a diversity of BMC-H subtypes (2, 30). Case in point, the BMC-H shell proteins from α- and ß-carboxysomes (CsoS1A and CcmK2, respectively) are structurally and phyletically distinct from one another, forming distant clades on a phylogenetic tree of all BMC-H protein sequences (2, 30). Therefore α- and ß-carboxysomes have distinct evolutionary histories, but converged on a functionally homologous BMC type (8, 9, 30). It is intriguing that α- and ß-McdB have seemingly also converged onto a similar mechanism of association with their respective carboxysomes; both mediated by a C-terminal invariant tryptophan.

Going forward, we will leverage this knowledge to examine conserved regions of CsoS1A and CcmK2 to identify co-occurring surface-exposed regions as candidate sites for interaction with the C-terminal motifs of α- and ß-McdB. Future studies such as this will help deepen our understanding of McdB localization to carboxysomes, and protein localization to BMCs in general.

### Tryptophan mediates the assembly of several viral and phage capsids

BMC shells share several analogous features to viral capsids (31). For instance, both are primarily comprised of hexameric proteins and some pentamers, the combination of which results in the characteristic polyhedral shape of BMCs and capsids (1, 32). Although structurally similar, BMC shells and viral capsids likely evolved independently, representing multiple convergent events (31). It is therefore insightful to compare these analogous structures and identify general features involved in self-assembly.

Intriguingly, many capsid proteins encode C-terminal tryptophan residues that are essential for the assembly of viral particles (33, 34, 35). Similarly, several carboxysome BMC-H proteins, including *Se* CcmK2, contain tryptophan residues at the interface of shell proteins that are often oriented toward the outer facet of the shell (36, 37). Albeit, none have been experimentally verified to be involved in assembly. However, the fact that tryptophan mediates interactions among viral capsid proteins and is found at the interface of BMC-H proteins suggests that tryptophan may be critical for mediating protein-protein interactions in these contexts. Whether McdB proteins interact with carboxysome shells via π–π stacking of tryptophan residues is an attractive mechanism of association for future study. Importantly, several groups are working to purify minimalized α- and ß-carboxysome shells to be used as tools in synthetic biology and biotechnological applications (38, 39). These minimalized shells could also serve as useful *in vitro* tools to reconstitute and study the molecular nature of McdB interactions with carboxysome components, and BMC-H proteins in particular.

### Kinesin-1 recognizes a tryptophan-acidic motif to interact with protein-based cargos

In eukaryotic cells, the motor protein kinesin-1 is critical for transporting diverse protein-based cargos on microtubules (40). How kinesin-1 recognizes and binds to a diverse set of cargo proteins to facilitate their transport has been under investigation for decades (41). It is now understood that these different cargo proteins all contain tryptophan-acidic motifs (such as EWD) that facilitate their binding to positively charged pockets on kinesin-1 (42).

Analogous to kinesin-1, McdB-like proteins must bind to diverse protein-based BMCs to facilitate their spatial regulation. Intriguingly, we show here that both α- and ß-McdBs tend to have acidic residues (often D) within 5 amino acids of the conserved tryptophan (see Figure 1). This is also true for the putative McdB-like proteins we identified for other BMCs (see Figure 4), although these contain aromatics residues other than tryptophan. It is therefore attractive to speculate that McdBs follow an analogous mechanism to bind BMCs as does kinesin-1 to its cargos, using tryptophan-acidic motifs to bind positively charged pockets on BMC shells.

Consistently, the carboxysome is known to have positively charged pockets within the pores of different shell proteins (43). Future investigations will therefore focus on the involvement of the conserved acidic residues in the C-termini of McdBs as well as the surface-exposed positive residues on BMC shells that could also mediate this association.

### The role of protein condensates in ParA/MinD-based positioning systems

McdB functions as an adaptor protein for the carboxysome positioning ATPase, McdA, which is a member of the ParA/MinD-family of positioning ATPases (13, 14, 19). This family is widespread in bacteria, and is responsible for spatially regulating a variety of genetic- and protein-based cargos (13, 14). The ATPases themselves do not interact directly with the cargo, but instead rely on adaptor proteins that either interact with, or are essential components of, the positioned cargos (15). How these adaptor proteins confer specificity to a wide variety of disparate cargos, ranging from the chromosome, divisome, flagella, chemotaxis arrays, and a diversity of BMCs, has been an active area of research for decades (13, 14, 15, 44).

Recently, the ability of some of these adaptor proteins to form condensates has been proposed to influence their localizations and functionality inside cells. For instance, the adaptor protein ParB which localizes to DNA molecules to aid in their segregation, has been shown to form condensates *in vitro* (45). ParB foci in the cell have also been seen to exhibit liquid-like behaviors (46), suggesting a dynamic, condensate-like mechanism underlying their formation and maintenance. Similar evidence exists for the co-complex of adaptor proteins, PomX and PomY, which localize cell division to mid-cell in some bacteria (47). The protein PomY forms liquid-like structures *in vivo* that, when perturbed, produce defects in cell division (48). Here we show a correlation between McdB localization to its carboxysome cargo, the ability of McdB to distribute carboxysomes, and its ability to form condensates both *in vitro* and *in vivo*. Whether the decrease in functionality we observed for our McdB variants is a direct consequence of the corresponding decrease in condensation activity will be an exciting area of future investigation.

## ACKNOWLEDGEMENTS

We would like to thank Dr. JK Nandakumar and Ritvija Agrawal for training and allowing us to use their SEC-MALS system. Dr. Joshua S. MacCready for providing access to previously curated lists of McdB protein sequences used here for the alignments. Claudia Mak for insightful conversations on the molecular nature of carboxysome shells and its contribution to our discussion here. And lastly, Dr. James Bardwell for allowing access to his CD spectrometer. This work was supported by the National Science Foundation to A.G.V. (NSF CAREER Award No. 1941966), Rackham Graduate Student Research Grant to J.L.B., and from research initiation funds provided by the MCDB Department to A.G.V.

## MATERIALS AVAILABILITY STATEMENT

Any materials used in this study are described in detail in Materials and Methods or can be accessed by contacting the authors.

## DATA AVAILABILITY STATEMENT

The data that supports the findings of this study are available in the supplementary material of this article or can be accessed by contacting the authors.

## MATERIALS AND METHODS

### Multiple sequence alignments

McdB amino acid sequences were obtained from our gene neighborhood analyses previously described for both α- (12) and ß-McdBs (11). Multiple sequence alignments (MSAs) were performed using Clustal Omega (49) and were exported and viewed using Geneious Prime (v 2020.02.02). Identity graphs were generated in Geneious Prime, and represent the percentage of pairwise residues that are identical in the alignment, including gap versus non-gap residues but excluding gap versus gap residues. Sequence logos were created using the above mentioned MSAs via WebLogo (v 2.8.2) (50).

### Construct design

All constructs used in this study were generated using Gibson Assembly (51). Cloning of plasmids was performed in chemically competent *E. coli* Top10 cells (Takara Bio). To replace native McdB with mutant variants in both *Se* and *Hn*, homology regions of 750 bp from both upstream and downstream of the native *mcdB* loci were added to the flanking regions of the generated constructs (52). Fluorescent fusions to proteins of interest were added to the indicated termini with a GSGSGS linker between the two proteins.

### Growth and transformation of Se strains

All *Se* strains were grown in BG-11 media (Sigma) buffered with 1 g/L HEPES, pH 8.3. Cultures were grown in a Minitron incubation system (Infors-HT) with 60 μmol m^−2^ s^−1^ continuous LED 5600 K light, 32°C, 2% CO_2_, and shaking at 130 RPM. Cells were transformed using 250-1000 ng of total plasmid DNA added to 300 µL of culture at OD_750_ = 0.7, and incubated in the dark for 16-24 hrs (52). Transformations were then plated on BG-11 media plus agar with the addition of 12.5 µg/mL kanamycin or 12.5 µg/mL chloramphenicol. Single colonies were picked and grown in BG-11 liquid media containing the same antibiotic concentrations, verified for full insertion via colony PCR, and then removed from antibiotics.

### Growth and transformation of Hn strains

All *Hn* strains were grown in ATCC® Medium 290: S6 medium for Thiobacilli (53). Cultures were grown in a Minitron incubation system (Infors-HT) at 30°C, 5% CO_2_, and shaking at 130 RPM. Competent *Hn* cells were generated by growing 1 L of log culture in 2.8 L flasks, which were harvested by centrifugation at 3,000g for 45 min. Cell pellets were washed twice with 0.5 volumes of ice-cold water, and finally resuspended in 1 mL of ice-cold water.

Competent cells were mixed with 250-1000 ng of total plasmid DNA and incubated on ice for 5 min. This mixture was then transferred to 5 mL of ice-cold S6 medium and incubated on ice for 5 min. Cells were then incubated for 16–24 hrs at 30°C, 5% CO_2_, and 130 RPM. Transformations were then plated on S6 media plus agar with 50 µg/mL kanamycin or 25 µg/mL chloramphenicol. Single colonies were picked and grown in S6 liquid media containing the same antibiotic concentrations, verified for full insertion via colony PCR, and then removed from antibiotics.

### Live cell fluorescence microscopy

For both *Se* and *Hn* cells, early log phase cultures grown in the absence of antibiotics were used for imaging. Two microliters of culture were spotted onto a 2 cm x 2 cm pad containing 1.5% UltraPure agarose (Invitrogen) + either BG-11 (for *Se*) or S6 media (for *Hn*). Cells were then imaged on a 35-mm glass-bottom dish (MatTek Life Sciences). All fluorescence and phase-contrast imaging was performed using a Nikon Ti2-E motorized inverted microscope controlled by NIS Elements software with a SOLA 365 LED light source, a ×100 objective lens (Oil CFI Plan Apochromat DM Lambda Series for Phase Contrast), and a Hamamatsu Orca-Flash 4.0 LTS camera. mNG constructs were imaged using a ‘YFP’ filter set (C-FL YFP, Hard Coat, High Signal-to-Noise, Zero Shift, excitation: 500/20 nm [490–510 nm], emission: 535/30 nm [520– 550 nm], dichroic mirror: 515 nm). mTQ constructs were imaged using a ‘CFP’ filter set (C-FL CFP, Hard Coat, High Signal-to-Noise, Zero Shift, excitation: 436/20 nm [426–446 nm], emission: 480/40 nm [460–500 nm], dichroic mirror: 455 nm).

### Image quantification using MicrobeJ

Image analysis including cell segmentation, quantification of foci spacing, and foci and cell intensities were performed using Fiji plugin MicrobeJ 5.13n (54, 55). Cell perimeter detection and segmentation were done using the rod-shaped descriptor with default threshold settings at a tolerance of 55 for both *Hn* and *Se* cells. Carboxysome foci were detected from both *Se* and *Hn* using maxima detection set to point detection with a tolerance of 1000 and the sharpen image filter selected. PCCs were calculated using ImageJ plugin JaCoP (56). Data were exported, further tabulated, graphed, and analyzed using GraphPad Prism 9.0.1 for macOS (GraphPad Software, San Diego, CA, https://www.graphpad.com).

### Protein expression and purification

Wild type and mutant variants for both *Se* and *Hn* McdB were expressed with an N-terminal His-SUMO tag off a pET11b vector in *E. coli* BL21-AI (Invitrogen). All cells were grown in LB + carbenicillin (100 µg/mL) at 37°C. One liter cultures used for expression were inoculated using overnight cultures at a 1:100 dilution. Cultures were grown to an OD_600_ of 0.5 and expression was induced using final concentrations of IPTG at 1 mM and L-arabinose at 0.2%. Cultures were grown for an additional 4 hours, pelleted, and stored at -80°C.

Pellets were resuspended in 30 mL lysis buffer [300 mM KCl; 50 mM Tris-HCl pH 8.4; 5 mM BME; 10% glycerol; 50 mg lysozyme (Thermo-Fischer); protease inhibitor tablet (Thermo-Fischer)] and sonicated with cycles of 10 seconds on, 20 seconds off at 50% power for 7 minutes. Lysates were clarified via centrifugation at 15,000 rcf for 30 minutes. Clarified lysates were passed through a 0.45 µm filter and loaded onto a 1 mL HisTrap HP (Cytiva) equilibrated in buffer A [300 mM KCl; 50 mM Tris-HCl pH 8.4; 5 mM BME; 10% glycerol]. Columns were washed with 5 column volumes of 5% buffer B [300 mM KCl; 20 mM Tris-HCl pH 8.4; 5 mM BME; 500 mM imidazole; 10% gylcerol]. Elution was performed using a 5-100% gradient of buffer B via an AKTA Pure system (Cytiva). Peak fractions were pooled and diluted with buffer A to a final imidazole concentration of < 100 mM. Ulp1 protease was added at 1:100 protease:sample, and incubated overnight at 25°C with gentle rocking. The pH was then adjusted to ∼10 and samples were concentrated to a volume of < 5 mL, passed through a 0.45 µm filter and passed over a sizing column (HiLoad 16/600 Superdex 200 pg; Cytiva) equilibrated in buffer C [300 mM KCl; 20 mM CAPS pH 10.2; 5 mM BME; 10% glycerol]. Peak fractions were pooled, concentrated, and stored at -80°C.

### SEC coupled to multi-angled light scattering (SEC-MALS)

For each sample analyzed, 50 µL at 1.5 mg/ml was passed over an SEC column (PROTEIN KW-804; Shodex) at a flow rate of 0.4 mL/min in buffer [150 mM KCl and 20 mM Tris-HCl, pH 8.2]. Following SEC, the samples were analyzed using an A_280_ UV detector (AKTA pure; Cytiva), the DAWN HELEOS-II MALS detector with an internal QELs (Wyatt Technology), and the Optilab T-rEX refractive index detector (Wyatt Technology). The data were analyzed to calculate mass using ASTRA 6 software (Wyatt Technology). Bovine serum albumin was used as the standard for calibration.

### Circular dichroism

For all protein samples analyzed, far-UV CD spectra were obtained using a J-1500 CD spectrometer (Jasco). All measurements were taken with 250 µL of protein at 0.25 mg/mL in 20 mM KPi, pH 8.0. Measurements were taken using a quartz cell with a path length of 0.1 cm. The spectra were acquired from 260 to 190 nm with a 0.1 nm interval, 50 nm/min scan speed, and at 25°C.

### Size-exclusion chromatography (SEC)

SEC was performed on full-length and truncated McdB proteins using a Superdex 75 Increase 10/300 GL (Cytiva) column connected to an AKTA pure system (Cytiva). 500 µL of sample at 1.5 mg/mL was passed through the column at 0.4 mL/min in buffer [150 mM KCl; 20 mM Tris-HCl pH 8.2] while monitoring absorbance at 220 nm.

### Microscopy of protein condensates

Samples for imaging were set up in 16 well CultureWells (Grace BioLabs). Wells were passivated by overnight incubation in 5% (w/v) Pluronic acid (Thermo-Fischer), and washed thoroughly with the corresponding buffer prior to use. All *Se* McdB samples were incubated for at least 30 minutes prior to imaging condensates, and all *Hn* McdB samples for at least 18 hours unless otherwise stated. Imaging of condensates was performed using a Nikon Ti2-E motorized inverted microscope (60 × DIC objective and DIC analyzer cube) controlled by NIS Elements software with a Transmitted LED Lamp house and a Photometrics Prime 95B Back-illuminated sCMOS Camera. Image analysis was performed using Fiji v 1.0.

### Quantification of phase separation via centrifugation

Centrifugation was used to quantify the degree to which McdB and its variants condensed under certain conditions, as previously described (57). Briefly, 50 µL of sample was incubated at the conditions specified for 30 minutes, and then centrifuged at 16,000g for 10 minutes at 20°C. The supernatant was removed and the pellet resuspended in an equal volume of McdB solubilization buffer [300 mM KCl, 20 mM CAPS pH 10.2]. Samples were then diluted into 4X Laemmli SDS-PAGE sample buffer. Pellet and supernatant fractions were visualized on a 4– 12% Bis-Tris NuPAGE gel (Invitrogen) by staining with InstantBlue Coomassie Stain (Abcam) for 1 hour and then destaining in water for 14-16 hours. The intensities of the bands were quantified using Fiji v 1.0 and resultant data graphed using GraphPad Prism 9.0.1 for macOS (GraphPad Software, San Diego, CA, www.graphpad.com).

### Expression of proteins to quantify condensate formation in E. coli

All *Se* McdB constructs were expressed as N-terminal mCherry fusions (58) on plasmids regulated by the pTrc promoter in *E.coli* MG1665. Overnight cultures were grown in 5 mL LB media + carbenicillin (100 μg/mL). The overnight culture was then diluted 1:50 into 5 mL AB Media + carbenicillin (100 μg/mL) supplemented with (0.2% glycerol; 10 μg/mL thiamine; 0.2% casein; 25 μg/mL uracil). *Hn* McdB was expressed as N-terminal mNG fusions (58) on plasmids regulated by the T7 promoter in *E.coli* BL21. Overnight cultures were grown in 5 mL LB media + carbenicillin (100 μg/mL). The overnight culture was then diluted 1:50 into 5 mL LB Media + carbenicillin (100 μg/mL). All cultures were allowed to grow at 37°C for until OD = 0.2-0.6 and then induced with 500 µM IPTG for *Se* strains, and 5 mM IPTG for *Hn*. The cultures continued to grow post-incubation for 3 hours before imaging. Cells used for imaging were prepared by spotting 2 µL of cells on to a 2% UltraPure agarose + AB medium pad on a Mantek dish. Images were taken using Nikon Ti2-E motorized inverted microscope controlled by NIS Elements software with a SOLA LED light source, a 100X Objective lens (Oil CFI Plan Apochromat DM Lambda Series for Phase Contrast), and a Hamamatsu Orca Flash 4.0 LT + sCMOS camera. mCherry signal was imaged using a “TexasRed” filter set (C-FL Texas Red, Hard Coat, High Signal-to-Noise, Zero Shift, Excitation:560/40 nm [540-580 nm], Emission: 630/75 nm [593-668 nm],Dichroic Mirror: 585 nm). For monitoring expression levels, cells were harvested either at the time of induction (t = 0 hours) or at the time of imaging (t = 3 hours). Cell lysates were normalized based on OD600, and were visualized via SDS-PAGE. Intensity of the bands corresponding to the McdB fusions were normalized to the background cell lysate and quantified. Image analysis was performed using Fiji v 1.0.

## SUPPLEMENTAL FIGURES

**Supplemental Figure S1:**
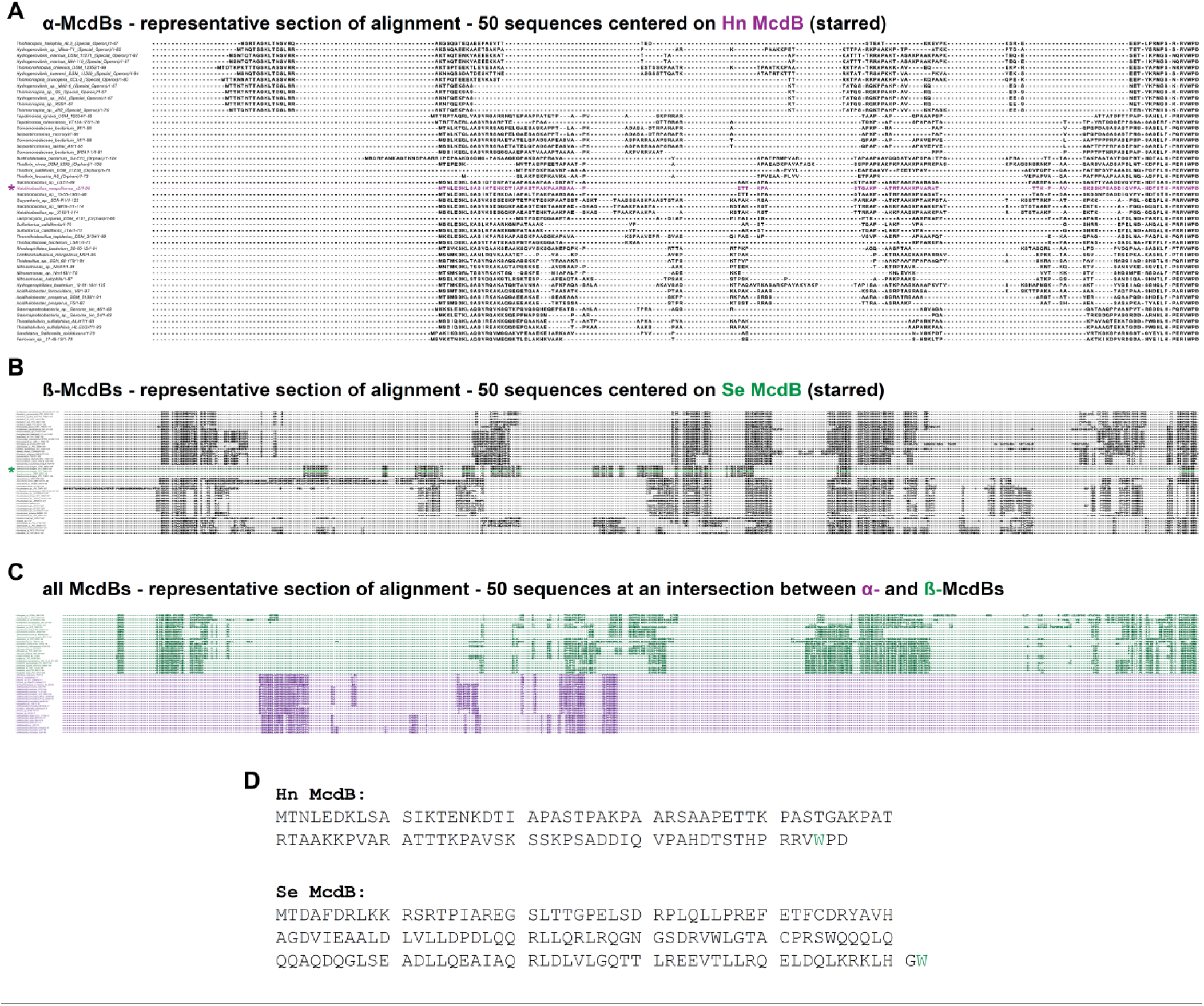
McdB amino acid sequences are highly variable. (**A**) A representative section of 50 sequences from the multiple sequence alignment of α-McdBs, centered on the sequence for *Hn* McdB (purple, starred). Dashes represent gaps in the alignment. (**B**) A representative section of 50 sequences from the multiple sequence alignment of ß-McdBs, centered on the sequence for *Se* McdB (green, starred). Dashes represent gaps in the alignment. (**C**) A representative section of 50 sequences from the multiple sequence alignment of all McdBs, centered on a region in which α-McdBs (purple) and ß-McdBs (green) align. Dashes represent gaps in the alignment. (**D**) Amino acid sequences of the full-length *Hn* and *Se* McdBs. The invariant tryptophan is colored green.

**Supplemental Figure S2:**
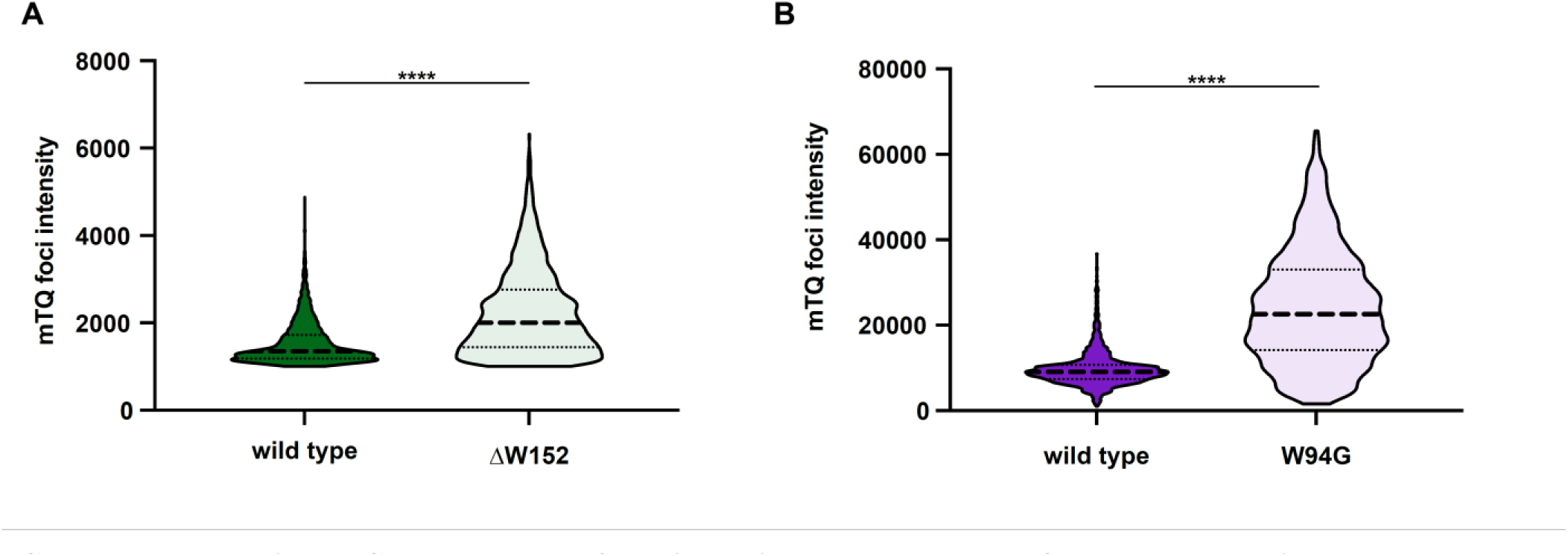
Removal of the invariant tryptophan of McdB results in carboxysome aggregation. (**A**) Quantification of RbcS-mTQ intensities from carboxysome foci in the indicated strains of *Se*. Graphs show medians and interquartile ranges from 3 biological replicates each with n > 1000 foci. **** p < 0.001 from Mann-Whitney U-test. (**B**) As in (A), but for *Hn* strains.

**Supplemental Figure S3:**
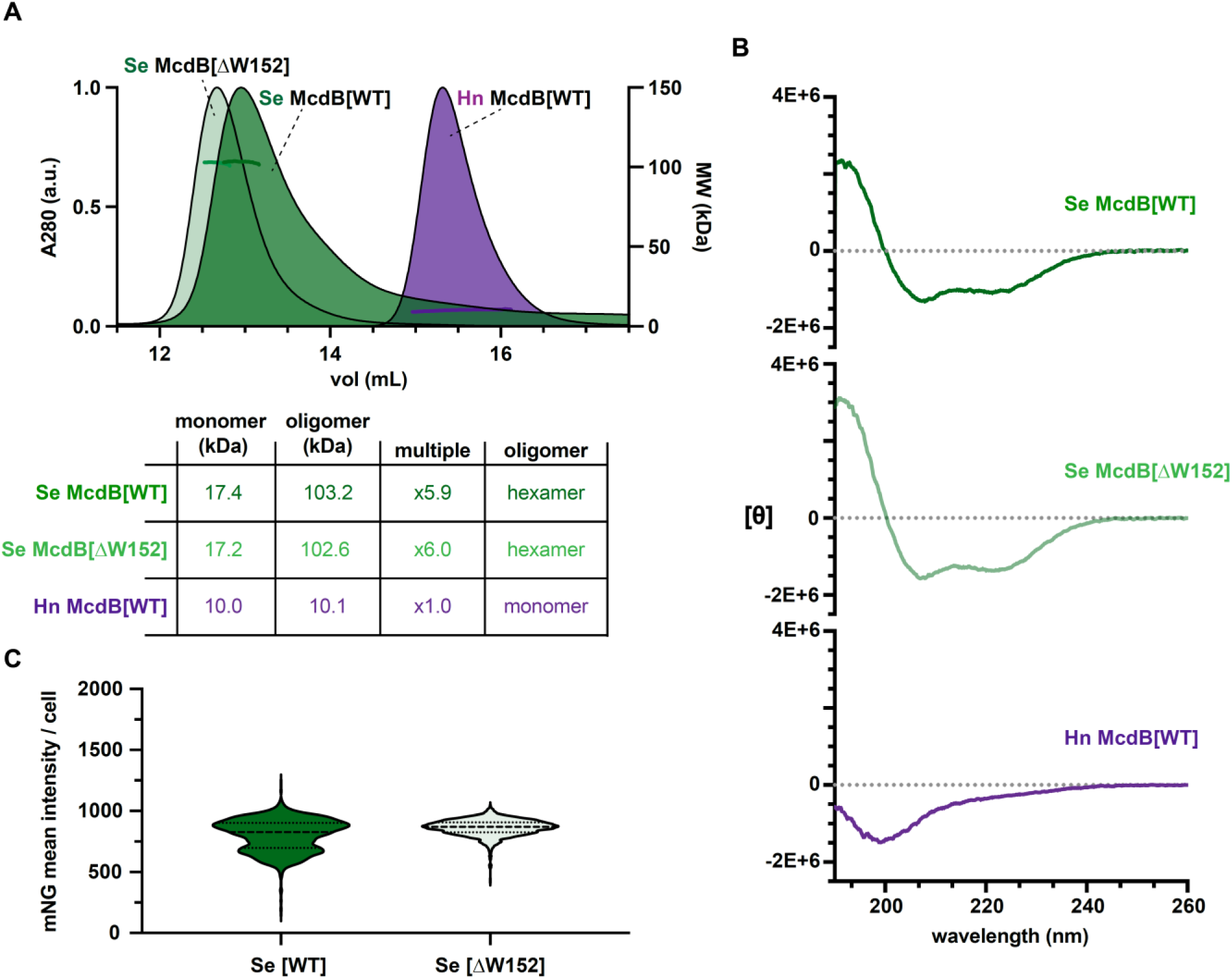
Mutations to the invariant Trp does not affect McdB protein stability *in vivo* or *in vitro*. (**A**) SEC-MALS for the indicated McdB proteins, with a summary table below. WT = wild type. (**B**) CD spectra for the indicated McdB proteins. (**C**) Quantification of mNG-McdB intensities per cell for the indicated strains. Graphs represent the medians and interquartile ranges for n > 500 cells for each strain.

**Supplemental Figure S4:**
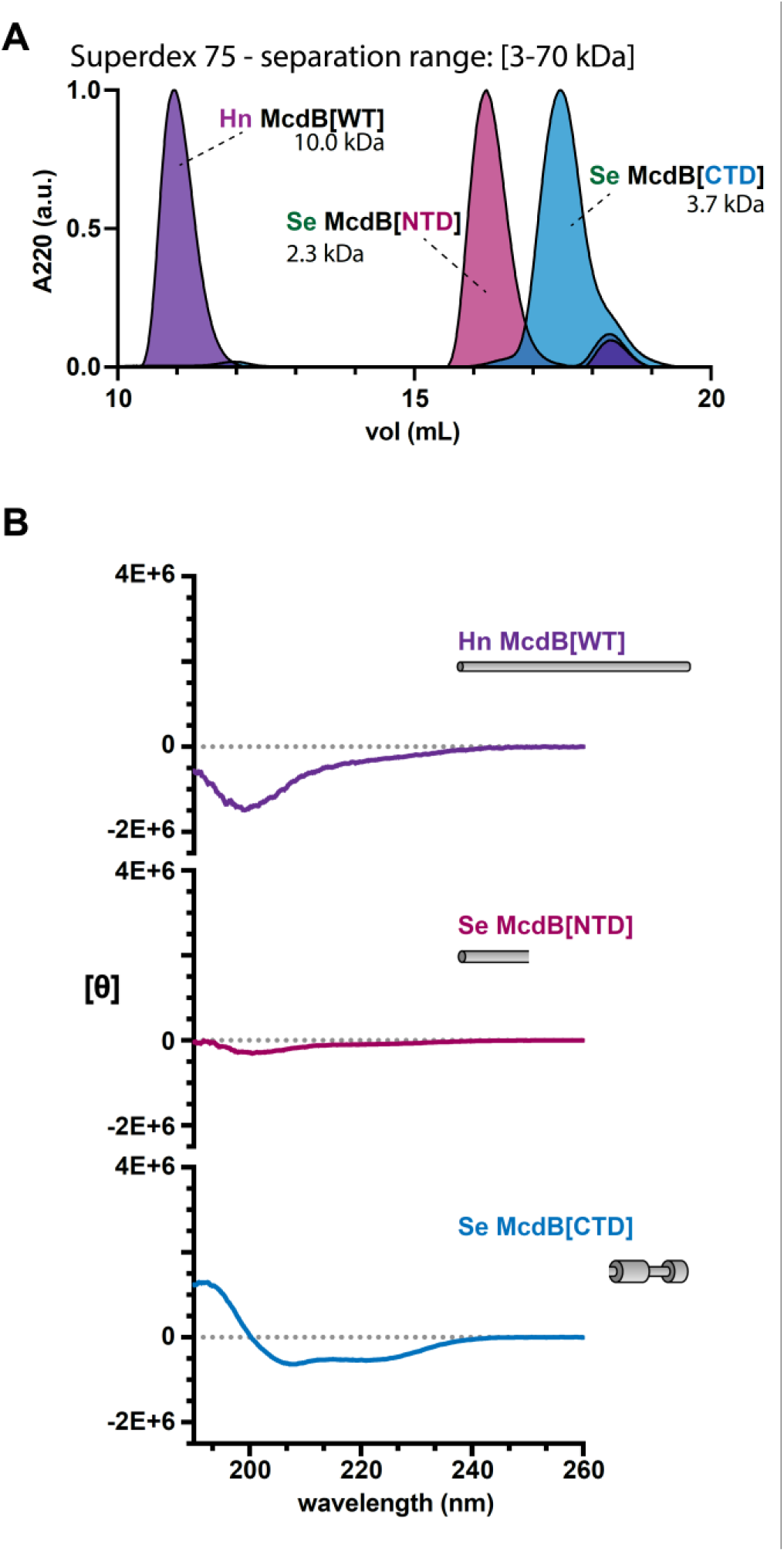
The C-terminus of *Se* McdB alone does not oligomerize, but remains α-helical. (**A**) SEC performed on the indicated column for the indicated protein variants. WT = wild type, NTD = N-terminal domain, CTD = C-terminal domain. Predicted monomeric weights are indicated for each construct. (**C**) CD spectra for the indicated proteins. Models of the constructs are shown, with wide cylinders representing α-helical regions and narrow cylinders representing region of intrinsic disorder.

**Supplemental Figure S5:**
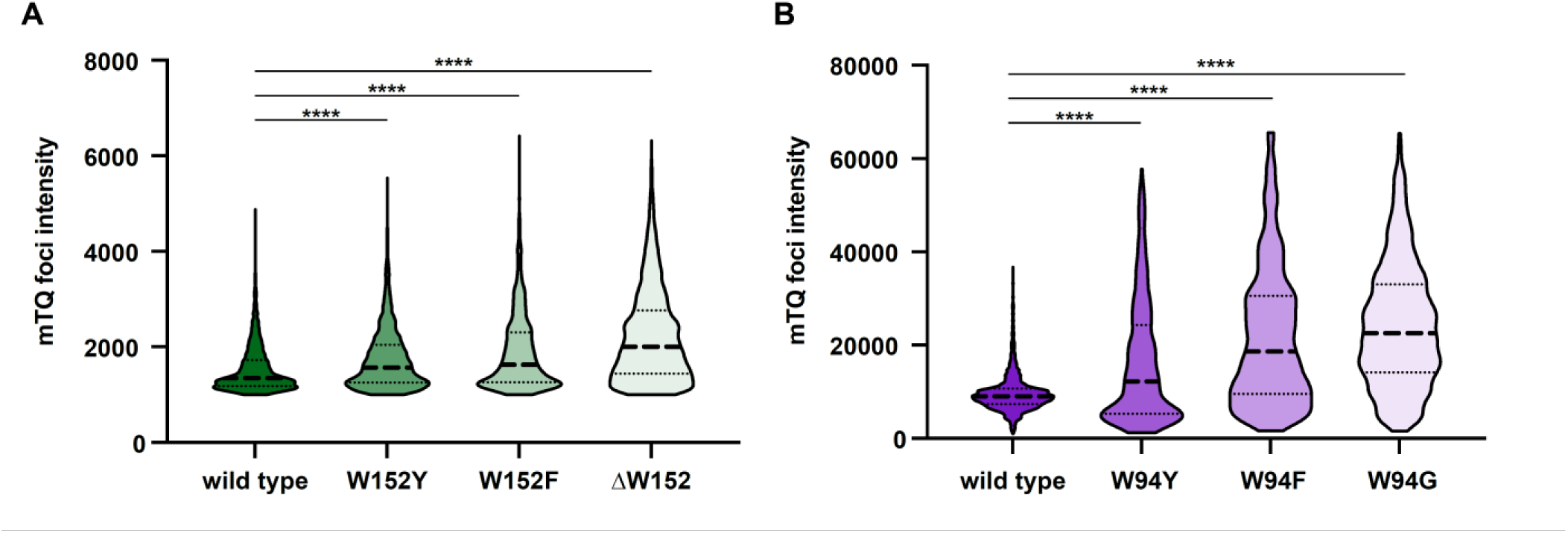
Changing the invariant tryptophan to other aromatic residues reveals a gradient of McdB function in positioning carboxysomes. (**A**) Quantification of RbcS-mTQ intensities from carboxysome foci in the indicated strains of *Se*. Graphs represent medians and interquartile ranges from 3 biological replicates each with n > 1000 foci. **** p < 0.001 from Mann-Whitney U-test. (**B**) As in (A), but for *Hn* strains.

**Supplemental Figure S6:**
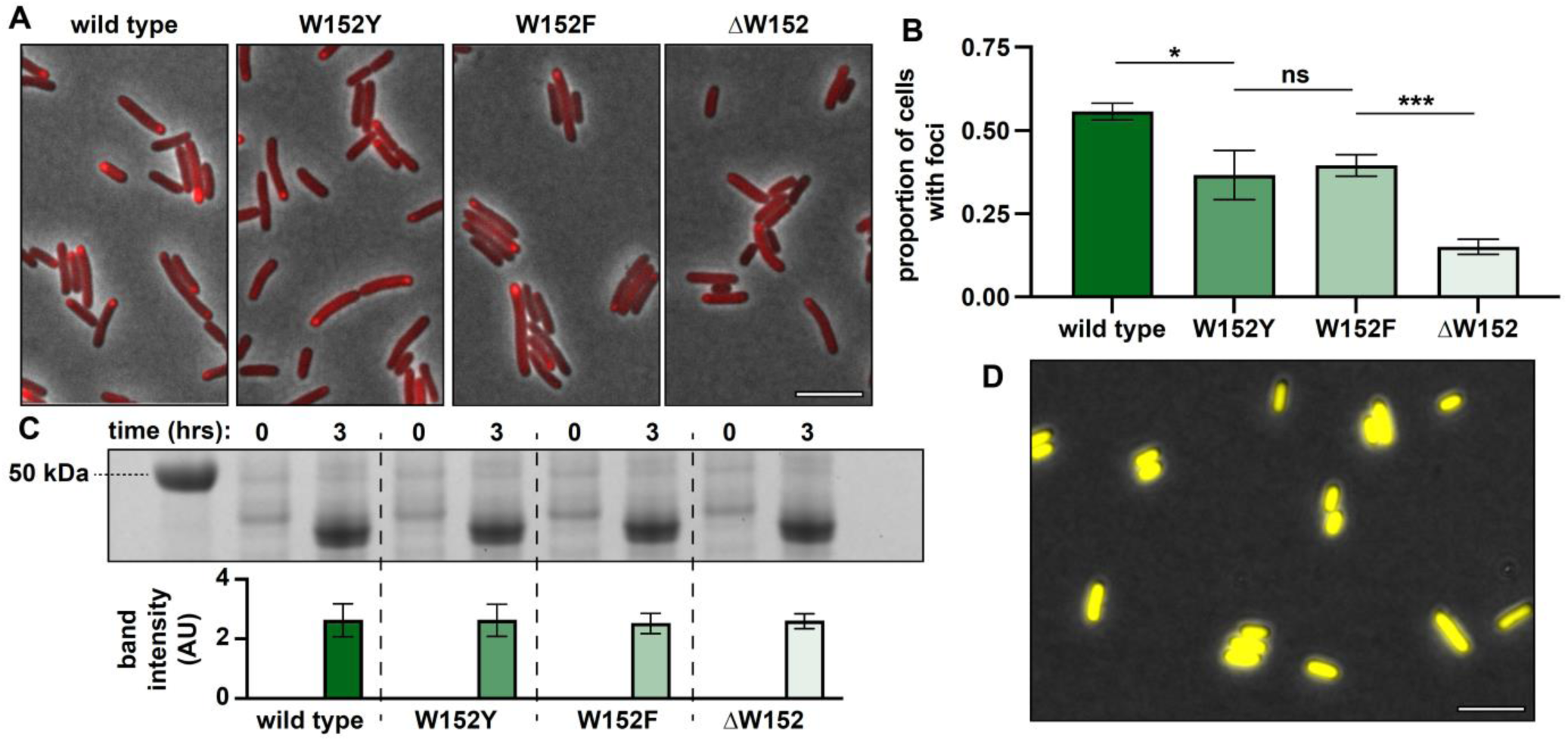
Differences in the solubilities of *Se* McdB aromatic substitutions in *E. coli*. (**A**) Representative microscopy image of the indicated *Se* mCherry-McdB variant after 3 hours of expression in *E. coli* MG1655. Phase contrast images are shown in black and white and overlaid with the fluorescence channel for mCherry-McdB as red. (**B**) Quantification of the proportion of cells from (A) with foci. Graphs represent means and standard deviations from 3 technical replicates each with n > 750 cells. *** p < 0.001, * p < 0.05, ns = non-significant from Welch’s t-test. (**C**) (*top*) SDS-PAGE gel from the experiment shown in (A). Cells were harvested at the indicated times of expression and standardized by OD600 prior to running on the gel. Expected size of mCherry-McdB constructs from *Se* is roughly 45 kDa. (*bottom*) Quantification of the normalized band intensities from the above gel. Graph represents means and standard deviations from 3 technical replicates. Comparisons of all variants at the 3-hour time point were non-significant from Welch’s t test. (**D**) Representative microscopy image of wild type *Hn* mNG-McdB after 3 hours of expression in *E. coli* BL21. Phase contrast images are shown in black and white and overlaid with the fluorescence channel for mNG-McdB as yellow. Note that the protein remained completely soluble, and so mutant variants of *Hn* McdB were excluded from further analysis. Scale bars are 5 µm and apply to all images.

